# ProteinConformers: large-scale and energetically profiled descriptions of protein conformational landscapes

**DOI:** 10.64898/2026.02.20.707011

**Authors:** Yihang Zhou, Chen Wei, Minghao Sun, Lin Wang, Jin Song, Fanding Xu, Yang Li, Wei Zheng, Yang Zhang

## Abstract

Modeling protein conformational landscapes is essential for understanding dynamics, allostery, and drug discovery, yet existing resources lack diverse conformational coverage, energetic annotations, or benchmarking standards. ProteinConformers (https://zhanggroup.org/ProteinConformers) provides 2.7 million geometry-optimized conformations generated with a multi-seed molecular dynamics strategy, paired with 13.7 million energy evaluations and 5.5 million similarity annotations. It delivers continuous landscapes from non-native to near-native states, benchmarking framework for multi-conformation generators, and an interactive analysis platform.

## Main

Understanding protein function requires capturing how structures dynamically interconvert across their conformational landscapes. Many proteins undergo allosteric and functionally critical transitions that occur at atomic resolution, and these motions are mostly governed by the underlying thermodynamic energy landscape^1-5^. Existing efforts to characterize protein conformational spaces fall into three broad strategies. Fragment-assembly methods construct decoy structures using Monte Carlo sampling^6-8^. Molecular dynamics (MD) simulations can generate conformations along trajectories initiated from experimentally resolved structures^9-13^. Ensemble extensions of deep learning based static structure predictors^14,15^ can generate multiple conformers. Despite these advances, current datasets and samplers exhibit major limitations: MD-based sampling is constrained by starting from native structures near the global energy minimum, existing conformer generators lack standardized benchmarks for assessing geometric plausibility and landscape-wide diversity, and available datasets provide limited energetic and structural similarity annotations, leaving the coupling between conformational variability and energetics insufficiently characterized.

To address these gaps, we developed ProteinConformers, a large-scale, energetically annotated resource of protein conformational landscapes. Using a multi-seed decoy sampling strategy that initiates MD simulations from hundreds of diverse starting conformations per protein, we generated over 2.7 million physically plausible structures spanning the full spectrum from non-native to near-native states, accompanied by 13.7 million energy evaluations and 5.5 million similarity annotations. ProteinConformers samples substantially broader conformational landscapes per protein than ATLAS database ^11^, while achieving atomic-level local plausibility comparable to the high-quality Top2018 dataset ^16^. Moreover, it enables systematic benchmarking of multi-conformer generators through our proposed evaluation framework based on the curated ProteinConformers-lite dataset. A user-friendly web platform supports querying, visualization, analysis, and data access for the community.

ProteinConformers offers an extensive characterization of structural and energetic landscape diversity across 734 proteins. The conformational ensembles span diverse conformation status (Fig. 1A) with a wide range of sequence lengths from 33 to 949 residues and an average of 247 residues (Fig. 1B-C). For each protein, we computed TM-score and RMSD between all sampled conformations and their native structures to quantify the global geometric plausibility and measure the non-native to near-native conformational spectrum (Fig. S1A). Energetic profiles were generated using five widely used statistical and physics-based energy functions, RW ^17^, RWplus^17^, EvoEF2^18^, Rosetta^19^, and FoldX^20^ (Fig. S1B). Structural classification using ECOD^21^ and CATH^22^ annotations further confirms that the dataset spans diverse protein fold families (Fig. S2).

**Figure 1.**
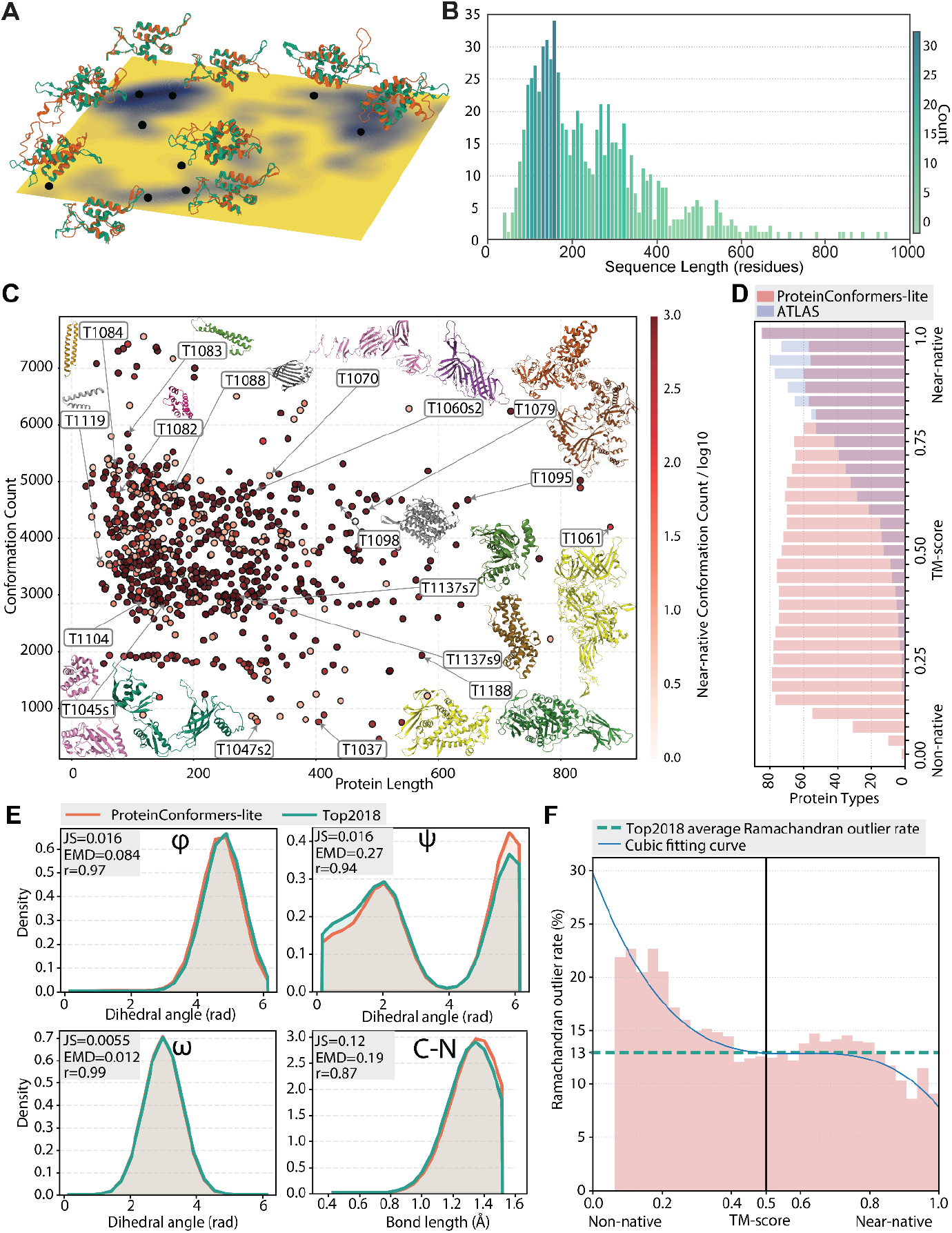
Overview and quality assessment of the ProteinConformers and ProteinConformers-lite datasets. (A) Schematic conformational energy landscape with representative conformers positioned in distinct minima of T1035, illustrating the breadth of physically realistic states captured by ProteinConformers. (B) Sequence-length distribution of ProteinConformers. (C) Conformer counts per protein, sorted by sequence length; points are colored by the log10 of near-native (TM-score ⩾ 0.5) conformation counts, with representative structures shown as insets. (D) TM-score coverage across proteins from ProteinConformers-lite and sampled ATLAS dataset, binned into 32 equal-width intervals from non-native to near-native. (E) Comparison of φ, ψ, ω dihedral angle and C–N bond length distributions between ProteinConformers-lite and Top2018 dataset, quantified by Jensen–Shannon divergence (JS), Earth Mover’s Distance (EMD), and Pearson correlation coefficient (r), demonstrating consistent local stereochemical quality. (F) Ramachandran outlier rates were averaged within 32 equal-width TM-score bins. The mean outlier rate of Top2018 dataset (13%, green dashed line) and the TM-score threshold of 0.5 separating non-native and near-native states are shown, with a cubic fit (blue curve) summarizing the trend.

We constructed ProteinConformers-lite, a curated benchmark subset of the ProteinConformers dataset containing 381,546 MD refined conformers across 87 CASP14 and CASP15 proteins (Fig. S3). These targets were selected because their seed decoys are generally higher quality and the underlying structures more challenging. Unlike previous benchmarks centered on near-native fluctuations around PDB structures, ProteinConformers-lite spans a broader conformational spectrum than the ATLAS dataset, with proteins distributed across non-native to near-native regions (Fig. 1D) and exhibiting a wider per-protein TM-score range (Fig. S4). Near native states were defined as those with TM-score greater than 0.5, a threshold indicating preservation of global fold integrity ^23^.

We evaluated the physical plausibility of ProteinConformers-lite by comparing its geometric properties with the high-quality Top2018 reference dataset. The distributions of dihedral angles and bond lengths in ProteinConformers-lite closely matched those in the Top2018 dataset (Fig. 1E), demonstrating that the local geometry of our benchmark achieves consistent stereochemical quality. Analysis of Ramachandran outlier rates revealed a consistent decrease as TM-score increased from non-native to near-native values, eventually falling below the average rate observed in the Top2018 dataset (13%). This trend indicates that near-native conformations in ProteinConformers-lite exhibit comparable stereochemical quality with Top2018 dataset (Fig. 1F).

A dual-axis benchmark framework was established to assess the diversity of the protein conformational landscape by calculating the coverage rate of the generated conformers to the benchmark dataset (Materials and Methods, Fig. S5), and to quantify residue-pair geometric statistics across distance and orientation features via Conformation Geometry Map (CGM) and its similarity score CGMS^cos^ and CGMS^mah^ (Materials and Methods, Fig. S6).

Then we systematically benchmarked five representative models 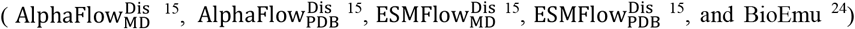 using our framework and ProteinConformers-lite dataset. Diversity analysis shows that BioEmu achieves the highest coverage under strict 5 kJ/mol thresholds, indicating effective sampling of low-energy regions, whereas distilled variants such as 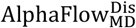 and 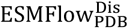 display consistently reduced coverage, reflecting narrower exploration around basins (Table S1). Plausibility of CGMS^cos^ emphasizes directional agreement in residue-pair statistics. The results show high similarity on distance features but substantial degradation on orientation features, indicating that current models generally recover distance distributions better than orientation statistics. CGMS^mah^ instead evaluates global geometric consistency by measuring Mahalanobis distances in the joint covariance space and converting them into similarity scores using a Gaussian kernel, resulting in more balanced assessment across feature types. BioEmu attains the highest scores on distance components under both metrics, reflecting stronger geometric modeling of inter-residue separations. Under CGMS^mah^, 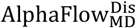 achieves scores comparable to BioEmu, suggesting similar overall geometric plausibility. In general, models fine-tuned on MD data (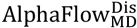 and 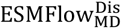) show modest improvements in CGMS^cos^ and comparable performance in CGMS^mah^ relative to their non–MD-tuned counterparts, indicating that MD refinements provide limited gains in local geometric realism (Table S2).

To enable efficient exploration of ProteinConformers, we developed an interactive web portal integrating query, visualization, and data access functions. The homepage provides a searchable table of all 734 proteins with multi field filters, linking to detailed dashboards for each target (Fig. 2A). Dashboards offer interactive 3D visualization with real time alignment between native and decoy structures (Fig. 2B), as well as native distance and rotation maps for global comparison (Fig. 2C). Each protein includes a fully annotated decoy table with geometric, similarity, energetic, and secondary structure metrics (Fig. 2D). Users can perform dynamic, in-browser analysis of the conformational landscape by filtering decoys based on structural similarity or energetic thresholds, with all statistical summaries and visualizations updating in real time. Flexible data export is supported through customizable metric selection (Fig. 2E), and one-click bulk downloads provide complete per-protein datasets with associated metadata (Fig. 2F). The portal enables rapid, interactive analysis without local computation.

**Figure 2.**
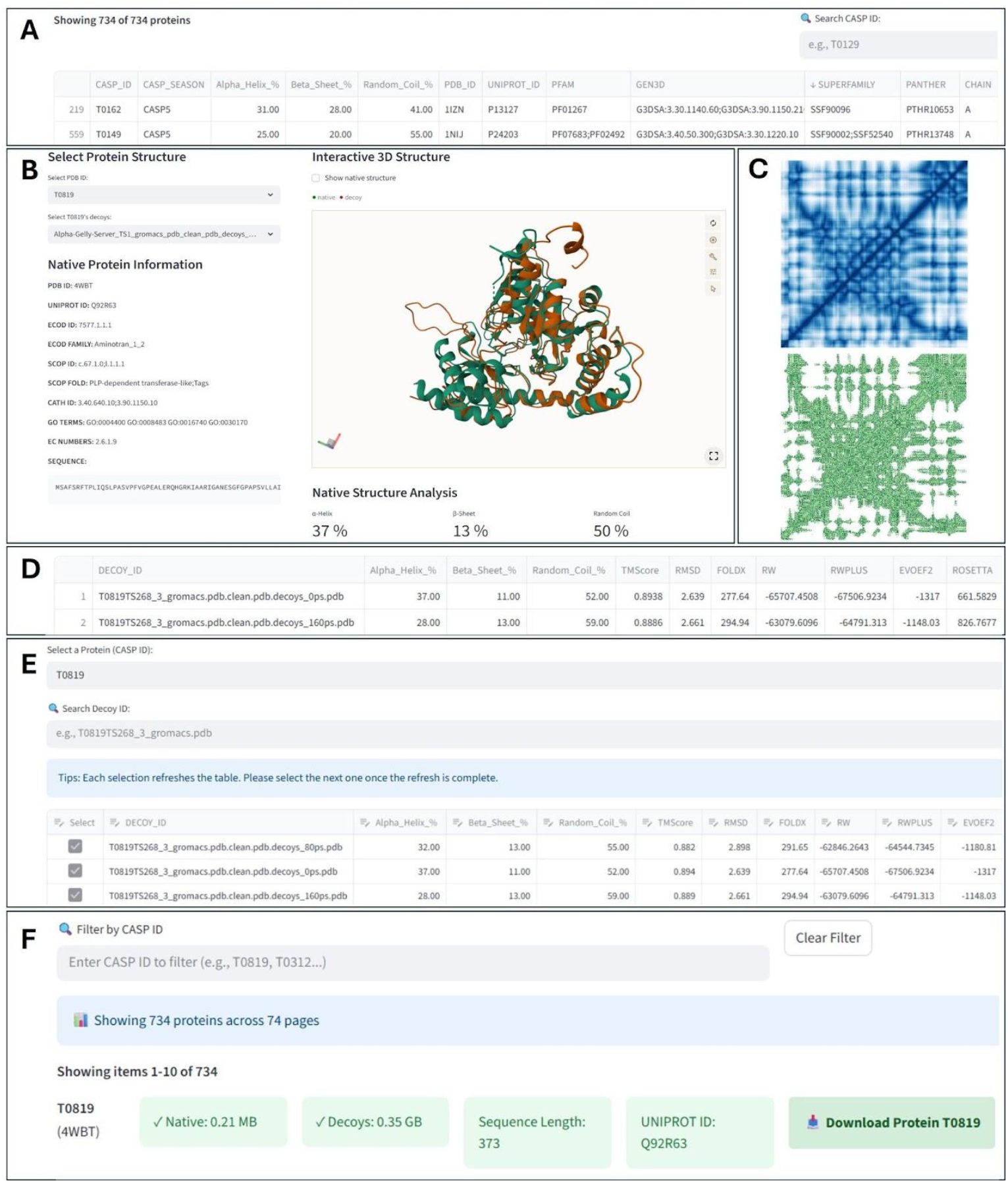
Interactive web interface of the ProteinConformers dataset. (A) Main overview table displaying all 734 proteins in the dataset, with sortable columns for structural metadata and CASP-related annotations. (B) Dashboard view for a selected target (e.g. T0819), showing an interactive 3D alignment of the native structure (green) with a selected decoy (orange), along with basic protein metadata and secondary structure composition. (C) Visualization of the native structure’s distance map (top) and orientation map (bottom), supporting global structural comparison. (D) Detailed table listing decoy models associated with the selected target, including secondary structure content, similarity scores, and energetic profiles. (E) Interface for selecting and filtering decoys by structural or energetic criteria. Selected decoys can be downloaded as a customized dataset. (F) Download panel summarizing metadata for the selected protein, including file sizes and sequence length, and providing options for downloading either the native structure or the full conformer set.

Taken together, ProteinConformers establishes a comprehensive, energetically annotated view of protein conformational landscapes, enabling systematic assessment of structural diversity and physical plausibility across modern multi-conformer generators. By integrating large-scale multi-seed sampling, detailed energetic profiling, and a unified evaluation framework, it provides a rigorous foundation for studying protein dynamics and allosteric mechanisms. Together with an interactive web platform for exploration and data access, ProteinConformers is positioned to support next-generation advances in protein conformation ensembles prediction, biomolecular modeling, and computational drug discovery.

## Materials and methods

### ProteinConformers Preparation

We manually downloaded all targets and corresponding predicted models (as decoys) submitted by participating groups worldwide from the Critical Assessment of protein Structure Prediction (CASP)^25^ across seasons 5 to 15. At the time of this data preparation, CASP16 predictions were not available. We developed and applied a comprehensive pipeline to remove redundant and erroneous structures and to complete incomplete decoys according to the following steps.

First, we manually retrieved, decompressed, matched, and initially filtered all sequence and structure files from the CASP repository. We then matched the sequence with structure filenames by string-based file name comparison. For structure files with identical names, we appended suffixes derived from their parent folders to distinguish. We then applied SeqKit^26^ to remove duplicate sequences. Finally, we manually inspected each pair. This process removed duplicate accessions, non-protein entries, and redundant sequences and associated predictions.

Second, we curated the structural files. Submissions from CASP teams frequently contained errors such as missing residues, incorrect residue ordering or numbering, missing atoms, unrecognized atom types, and erroneous bonding. Similar issues were also observed in the native PDB files provided by CASP, together with inconsistencies between the reference sequence files and the sequences extracted from the native structures, which is often due to limited resolution in certain regions of the experimentally determined structures. To address these issues, we used the sequence files as the reference and performed a clean process. For each prediction, chains were extracted from structure files using Biopython ^27^ and aligned to the sequence reference. The alignments were classified into three mutually exclusive categories: *same* (full-length with identical residue order, for example, reference sequence ABCD vs. extracted sequence ABCD), *disorder* (a contiguous subsequence of the reference caused by order-preserved internal or terminal deletions, for example, reference ABCD vs. extracted ABD), and *mismatch* (any substitution, insertion, or reordered segment, for example, reference ABCD vs. BACD or ABED). Missing atoms were rebuilt with OpenBabel ^28^, while structures with unrecognized atom types or bonding errors were excluded. Oligomeric or hetero-complex models were excluded from the dataset. Based on the categories from the previous step, the *same* models were accepted without modification. The *disorder* models were retained only if their residue numbers matched the reference structure; otherwise, the coordinates would be automatically renumbered to restore one-to-one correspondence. All *mismatch* entries were discarded.

Third, when multiple experimental native structures were available for a target, we manually selected a single representative. Selection criteria included secondary validation in the PDB, sequence length, and structural resolution.

Fourth, additional filtering was performed based on data availability. A target was included in the benchmark only if at least one native structure and 100 decoys were available. For targets below this threshold, additional decoys would be generated using 3DRobot ^8^. Targets that failed to obtain sufficient decoys were removed.

After these steps, all native structures and decoys underwent further physics-based optimization through MD simulations. Structures that crashed during MD process were abandoned.

### Molecular Dynamics Protocol

We developed and applied a full-atom molecular dynamics simulation pipeline to optimize all conformations at atomic resolution, and all conformations that failed to converge during this process were excluded.

Atomic MD simulations are performed using GROMACS 2023 ^29^. Each protein conformer follows the same workflow. Topology construction and solvation: The OPLS-AA force field is used for topology generation, together with the TIP3P water model. Each protein is centered in a dodecahedral box, and the box is filled with pre-equilibrated SPC216 water. Na^+^ and Cl^−^ ions are added to neutralize the total charge. Steepest-descent minimization is applied until the largest force on any atom falls below 1000 kJ mol^−1^nm^−1^ (maximum 50,000 steps), thereby minimizing the energy, eliminating steric clashes and unrealistic geometries. The NVT phase (100 ps, 300 K) employed the V-rescale thermostat with positional restraints on heavy atoms to stabilize temperature. The subsequent NPT phase (100 ps, 1 bar) uses the Parrinello-Rahman barostat after partially releasing restraints to equilibrate density. Periodic-boundary artifacts are eliminated by recentering and re-imaging the trajectory. Restraints are removed, and a simulation between 125 ps to 375 ps is executed for simulation. Snapshots are extracted every 25 ps.

All the simulations are executed on high-performance computers equipped with AMD EPYC 7763 (64-core, 2.45 GHz) processors.

### Energetic and Similarity Profiles

We annotated the energetic profile for each conformation using RW ^17^, RWplus ^17^, EvoEF2 ^18^, Rosetta (version: REF2015) ^19^, and FoldX (version: 20241231) ^20^.

We calculated pairwise conformational similarity metrics between each decoy to the native structure, including TM-score ^23,30,31^ and RMSD ^32^.

### ProteinConformers Dataset statistics

ProteinConformers includes 734 native structures and 2,750,261 optimized atomistic conformations generated from 352,146 seed decoys. On average, each protein runs MD simulation from 480 different initial structures and has 3,747 conformations. Each conformation has 5 energetic profiles and 2 similarity metrics, which in total result in 13,751,305 energetic profiles and 5,501,990 similarity annotations. The 734 distinct proteins range in sequence length from 33 to 949 residues, and an average length of 247 residues. The aggregate cost is approximately 40 million CPU hours.

### Benchmark Dataset

To highlight the capabilities of the ProteinConformer dataset and to facilitate systematic evaluation of existing protein conformation generative models, we developed a compact benchmarking suite, referred to as ProteinConformers-lite. The conformational landscapes in this dataset were collected from ProteinConformers, including conformers from CASP14 and CASP15. We selected the most recent two CASP seasons because their seed decoys are generally of higher quality and the corresponding targets are overall more challenging.

The ProteinConformers-lite benchmark dataset comprises 87 proteins with an average sequence length of 305 residues, a median of 255 residues, and a maximum length of 949 residues (Fig. S1). These 87 proteins include 40,387 seed decoys, 381,546 conformers, and 1,907,730 energetic annotations.

We depolyed 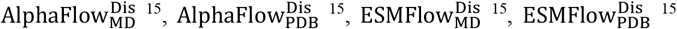, and BioEmu ^24^ followed their official tutorials. For every software, we generated 3,000 conformations for each protein in ProteinConformers-lite, and assessed the diversity and plausibility of the generation using our metrics.

### Top2018 dataset comparison

The Top2018 dataset is a high-quality curated reference collection of protein residues for plausibility reference ^16^. We downloaded the complete Top2018 dataset, which contains 8,307 high-quality, low-redundancy protein chains. Ramachandran outlier rates were computed using PyRosetta ^33^ with the *Ramachandran* scoring function, where *phipsi_in_forbidden_rama* was applied to determine whether each residue dihedral angle falls into a disallowed region, excluding terminal residues. For bond length and bond angle statistics, we used Biopython ^27^ to calculate arithmetic means of the main chain bond lengths. Dihedral angles were first mapped onto the [0, 2π) interval and then averaged using the circular mean, which include φ (C_(i-1)_-N_(i)_-Cα_(i)_-C_(i)_), ψ (N_(i)_-Cα_(i)_-C_(i)_-N_(i+1)_), and ω (Cα_(i)_-C_(i)_-N_(i+1)_-Cα_(i+1)_).

### ATLAS dataset comparison

The ATLAS dataset provides MD trajectories generated from a single initial conformation per protein. To enable a direct comparison of conformational diversity between ATLAS and ProteinConformers, we randomly selected 87 proteins from ATLAS (Table S3) as the same number of proteins in ProteinConformers-lite. For each target, trajectory files were downloaded and conformations were extracted following the official ATLAS processing protocol, yielding 30,000 conformations per protein. TM-scores were computed using the corresponding processed native structures as references.

### Diversity Evaluation Metrics

To quantitatively benchmark generative models against this reference set, we adopt an energy-threshold-based overlap analysis over discretized 2D free energy landscapes. In this study, we adopt the dimensionality reduction technique Principal Component Analysis (PCA) to project the high-dimensional conformational data onto a low-dimensional space. Each conformer’s energy is projected into a reduced-dimensional free energy landscape via PCA, followed by Boltzmann-weighting to estimate the free energy of each state on this landscape by weighting the contribution of each conformation according to its probability as determined by the Boltzmann distribution. For a given energy threshold *τ*, which is a cutoff value used to determine which protein structures (conformations) are considered stable and realistic enough to be included in the analysis, we evaluate three complementary metrics between a reference ensemble *A* (ProteinConformers-lite) and a generated ensemble *B*, including *Interaction, Coverage*, and *Jaccard Index*.

*Interaction* measures the absolute number of shared low-energy regions:

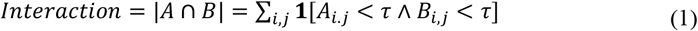

where *i, j* are the indices of the grid of the Boltzmann-weighted estimation of the energy landscape.

*Coverage* indicates the fraction of ProteinConformers-lite’s low-energy regions recovered by the generative model:

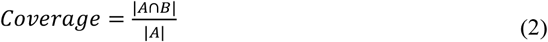

*Jaccard Index* provides a symmetric measure of shared low-energy regions:

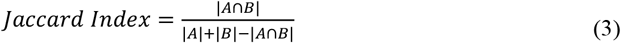

These metrics are evaluated under increasingly relaxed energy thresholds (5, 10, 20 kJ/mol), allowing assessment of both strict and broader sampling capabilities. The detailed comparison results are provided in Table S1.

### Plausibility Evaluation Metrics

To evaluate the realism of generated ensembles, we propose the Conformation Geometry Map (CGM) and its derived Conformation Geometry Map Similarity (CGMS).

#### CGM Construction

Given a protein of length *N* and an ensemble of *M* conformations, 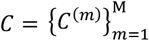, we compute four inter-residue geometric features *G* ∈ [*D, Ω, Θ, Φ*] for each conformation, following definitions from trRosetta ^34,35^ and DeepPotential ^36^ (Table 1).

**Table 1.** Geometric features used for CGM. For Ω, Θ, and Φ, a pseudo *C_β_* is used for glycine.

| Symbol | Type | Atoms Involved | Symmetry | Description |
| --- | --- | --- | --- | --- |
| $D$ | Euclidean Distance | $C_{\alpha,i}, C_{\alpha,j}$ | Symmetric | Distance ($D_{ij} = D_{ji}$) |
| $\Omega$ | Dihedral Angle | $C_{\alpha,i}, C_{\beta,i}, C_{\beta,j}, C_{\alpha,j}$ | Symmetric | Rotation about $C_{\beta,i} - C_{\beta,j}$ virtual axis |
| $\Theta$ | Dihedral Angle | $N_i, C_{\alpha,i}, C_{\beta,i}, C_{\beta,j}$ | Asymmetric | Orientation of $C_{\beta,j}$ to residue $i$ backbone |
| $\Phi$ | Planar Angle | $C_{\alpha,i}, C_{\beta,i}, C_{\beta,j}$ | Asymmetric | Position of $C_{\beta,j}$ to $C_{\alpha,i} - C_{\beta,i}$ bond |

For each residue pair (*i, j*), we calculate the mean (*μ*), standard deviation (*σ*), and skewness (*γ*) of each geometric feature across the ensemble. The calculation of angle features considers circularity. The resulting feature tensor is *P*^*G*^ ∈ *R*^3×*N*×*N*^, and the full CGM is the concatenation *P* = [*P*^*D*^, *P*^*Ω*^, *P*^*Θ*^, *P*^*Φ*^] ∈ *R*^12×*N*×*N*^. The Euclidean distance 20Å is used as the cutoff to filter the residue pair for CGMS.

#### CGMS Computation

Computation involves two variants: CGMS^cos^ and CGMS^mah^. For a given test dataset and the reference ProteinConformers-lite ensemble, the respective CGM matrices, denoted as *CGM*^*Test*^ and *CGM*^*Ref*^ are computed under the condition that the test dataset samples the same set of proteins as the reference. For each valid residue pair (i, j), a three-dimensional vector *u* = (*μ, σ, γ*) is extracted from both maps.

For CGMS^cos^, the average cosine similarity is calculated to evaluate the overall directional correspondence between the descriptor vectors, with less sensitivity to local deviations. The metric is defined as the mean similarity over all aligned residue pairs:

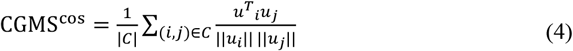

where *C* denotes the set of all valid residue pairs.

CGMS^mah^ incorporates both directional and magnitude alignment through a composite Mahalanobis distance and a Gaussian radial basis function, imposing a stronger penalty on local discrepancies. The Mahalanobis distance is defined as:

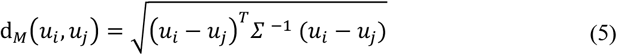

where *Σ* in (5) is the covariance matrix between two vectors. This distance is then transformed via a Gaussian kernel to compute the similarity:

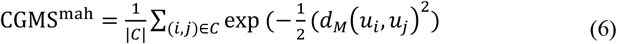

Higher CGMS indicates better agreement with the native ensemble’s inter-residue geometric statistics. The detailed comparison results are provided in Table S2.

### Website implementation

The ProteinConformers portal is implemented as an interactive data-driven web application built with Streamlit (v1.32), an open-source Python framework for rapid development of scientific dashboards. The Streamlit service is deployed on a dedicated backend and is embedded within the main site through HTML, allowing seamless integration of dynamic content without coupling the application to the primary presentation layer. The enclosing HTML page is delivered by an Apache HTTP Server, thereby decoupling the interactive visualization engine from the static web infrastructure while preserving performance and security best practices.

The application architecture incorporates a multi-tab layout, implemented using the streamlit-option-menu library, which allows users to navigate between sections such as Home, Data Table, Structure Browser, Download Center, and About. Each section is backed by efficient data caching via @st.cache_data decorators, which reduce redundant I/O and enable fast stateful browsing across user sessions. The Home tab offers summary metrics and structural composition plots; the Table and Download tabs enable rich, multi-parameter filtering of over 2.7 million conformations using cross-linked annotations (CATH, SCOPe, ECOD, GO terms, EC numbers, etc.); while the Browser tab features an interactive 3D visualization module powered by the streamlit-molstar extension ^37^ for PDB file rendering, and it providing users with an interactive tool to inspect, rotate, and analyze protein conformers in detail. The interactive 2D charts, such as scatter plots and heatmaps, are rendered using the Plotly ^38^, allowing users to dynamically explore the data.

The portal supports real-time filtering through a combination of pandas ^39^ operations and Streamlit widgets, with built-in pagination to handle large tables and performance-optimized rendering for charts. Extensive custom CSS is injected to declutter the UI and suppress non-essential components such as default toolbars and deploy buttons, enhancing the user experience. Additionally, protein-specific decoy ensembles, native structures, and energy landscapes are integrated on demand through efficient file I/O and optional download modules, which estimate data size and dynamically build user-selected archives. The entire application is deployed on a web server to ensure stable and public accessibility for the research community.

## Data Availability

All processed data in ProteinConformers is freely accessible at https://zhanggroup.org/ProteinConformers without registration requirements. The benchmark dataset ProteinConformers-lite is also available at https://huggingface.co/datasets/Jim990908/ProteinConformers/tree/main. The benchmark codes and instructions are available at https://github.com/auroua/ProteinConformers.

## Authors’ contribution

Ya.Z. contributed to the conception and design of the study; Yi.Z. generated the dataset; C.W., M.S., J.S. and F.X developed the website. Yi.Z., C.W., L.W., M.S., J.S. designed the methodology; C.W., M.S., L.W., J.S., F.X., Y.L., W.Z. contributed to the discussion and experiment design; Yi.Z., C.W., M.S., L.W., J.S., F.X., Y.L., Ya.Z. wrote the paper. All authors proofread and approved the final manuscript.

## Supplementary data

Figure S1. Example of similarity score and energetic annotation distributions in ProteinConformers

Figure S2. Distribution of fold classes in ProteinConformers

Figure S3. The 3D native structures of all 87 proteins in ProteinConformers-lite

Figure S4. Violin plot of TM-score range per protein

Figure S5. Example of free energy landscapes comparison from ProteinConformers-lite and generative models

Figure S6. Illustration of CGM and CGMS

Table S1. Diversity benchmark on ProteinConformers-lite under different energy thresholds

Table S2. Plausibility benchmark on ProteinConformers-lite

Table S3. IDs of sampled proteins from ATLAS

